# Lcn2 protects from cisplatin induced AKI by regulating DNA damage response via EGFR activation

**DOI:** 10.1101/2024.05.03.592429

**Authors:** Aniela Zablocki, Eloise Marques, Lucie Yammine, Clément Nguyen, Fabiola Terzi, Morgan Gallazzini

## Abstract

**Background:** Lipocalin 2 (Lcn2), a small-secreted protein, is an established sensitive biomarker of kidney injury. In Cisplatin (CDDP) induced acute kidney injury (AKI), Lcn2 expression is swiftly and strongly increased in suffering renal medullary tubules. While recent reports correlates Lcn2 expression in cancer cells with CDDP resistance, the role of Lcn2 in kidney tubule damaged by CDDP remains unknown.

**Methods:** To better understand the role of Lcn2 in CDDP-induced AKI, experiments on *Lcn2^+/+^* or *Lcn2^−/−^* mice as well as immortalized kidney cells knock-down (KD) for Lcn2 were conducted. Kidney function and injury were assessed using standard techniques. Cellular and molecular mechanisms were studied in WT and Lcn2 KD cells in combination with pathways inhibitors in order to gain insight Lcn2 driven mechanisms.

**Results:** In animal injected with CDDP, Lcn2 was upregulated mostly in kidney inner medulla collecting duct while it was reabsorbed in the proximal tubules. *Lcn2^−/−^* in mice significantly decreased kidney function compare to WT mice, while it increased parenchymal damage due to increased cell death and cast formation. Interestingly, while little to no damage were present in the medulla of CDDP injected WT animal, a clear increase of medulla tubular lesions was observed in *Lcn2^−/−^* mice. Using Inner Medullary Collecting Duct cells (mIMDC-3), we showed that Lcn2 KD induces a p53-dependent apoptosis upon Cisplatin exposure while no effect on necrosis was observed. Finally, we demonstrated that Lcn2 drives CDDP resistance to apoptosis through the activation of the pro-survival EGFR pathway.

**Conclusions:** We established that Lcn2 is a renoprotective protein in CDDP-induced AKI via the activation of EGFR pathway. This new mechanism might represent a new approach for the treatment of AKI.

## INTRODUCTION

Cisplatin (CDDP) is a platinum-based chemotherapeutic agent commonly used for many types of cancer such as lung, testicular, ovarian and other cancers (Wang & Lippard, 2005 ; Siddik, 2003). However, many side effects affecting healthy organs, mainly the kidney, limit its clinical application. In fact, a third of patients on treatment will develop Acute Kidney Injury (AKI) (Arany & Safirstein, 2003). AKI is a major kidney disease characterized by the rapid loss of renal function and is associated with high rates of morbidity and mortality. Furthermore, cisplatin-induced AKI is contributing to the development of chronic kidney disease (CKD) (Shi *et al*, 2018 ; Fu *et al*, 2019) The mechanisms of cisplatin-induced AKI are complex and include oxidative stress, DNA damage, inflammation and also proximal and distal tubular cell death by apoptosis and necrosis (Miller *et al*, 2010 ; Pabla & Dong, 2008 ; Wei *et al*, 2007). However, the molecular and cellular pathways associated with these phenotypes are not fully characterized. Therefore, a better understanding of the cisplatin-induced AKI mechanisms is needed in order to find a novel renoprotective therapy.

Lipocalin-2 (Lcn2, also known as neutrophil gelatinase-associated lipocalin (NGAL) or 24P3), is a small secreted protein belonging to the lipocalin superfamily (Flower *et al*, 1991). Lcn2 was originally described as a transporter of small hydrophobic molecules and could also act as an anti-bacterial agent by binding the siderophore-iron complex essential for bacterial proliferation (Chakraborty *et al*, 2012). In addition to these initially described roles, Lcn2 appears to be involved in many physiological processes such as cancer development and cellular stress. Some studies have reported Lcn2 as a protein with antioxidant properties (Bahmani *et al*, 2010 ; Roudkenar *et al*, 2008), but also as being pro- or anti-apoptotic depending on the context (Kehrer, 2010). However, the molecular mechanisms underlying the role of Lcn2 in these contexts are not yet defined.

During AKI, Lcn2 has been shown to be highly induced in the kidney both in humans and in mouse models (Grigoryev *et al*, 2013). Due to its rapid increase in urine and plasma as early as 2 hours of AKI, Lcn2 is considered as a biomarker of injury status (Mishra *et al*, 2003). In line with this, Paragas et al. (Paragas *et al*, 2011) demonstrated that Lcn2 expression is induced in Thick Ascending Limbs of Henle (TAL) and the collecting ducts (CD) in the outer stripe of the inner medulla. Furthermore, they reported that the damaged nephrons were the source of Lcn2. In this study, no mRNA expression of Lcn2 was observed in the proximal tubule. Hence, Lcn2 may be a cell autonomous stress response protein important for the TAL and CD integrity upon injury. Unfortunately, the mechanisms by which Lcn2 exerts its nephro-protective action in the context of cisplatin have not been described. Nevertheless, it has been recently reported in endometrial carcinoma cells, that Lcn2 expression was increasing cell survival upon Cisplatin treatment (Miyamoto *et al*, 2016a). We therefore decide to investigate, in a renal context, the cellular and molecular mechanisms Lcn2 regulates and its impact on tubular injury during AKI.

Here, we demonstrate that cisplatin-injected mice invalidated for Lcn2 expression develop more parenchymal lesions than wild-type mice. These effects were consistent *in vivo* and *in vitro* with an increase in apoptosis induced by cisplatin in the absence of Lcn2. *In vitro*, Lcn2 invalidation led to a global alteration of the DDR response in renal tubular cells, causing an increased apoptosis. Finally, we demonstrate that Lcn2 mediates its pro-survival effect via the activation of the Epidermal Growth Factor Receptor (EGFR). Together, our findings describe the mechanism by which Lcn2 protects the kidney from genotoxic stress.

## RESULTS

### Lcn2 prevent Cisplatin-induced Acute Kidney Injury

In order to elucidate the role of Lcn2 in CDDP-induced AKI, we first characterized Lcn2 expression pattern in kidney tubular cells up to 4 days after Cisplatin injection. Indeed, while Lcn2 expression in kidney tubule after ischemic injury is well described, the effect of Cisplatin on Lcn2 expression in kidney parenchyma remains elusive. With this mind, *Lcn2^+/+^* and *Lcn2^−/−^*mice were sacrificed 1-, 2- or 4-days after cisplatin injection and Lcn2 protein kidney localization was monitored by immuno-fluorescence in tissue sections. While Lcn2 expression was detected in the inner medulla in control animals (Figure 1A and supplementary Figure 1A), a progressive induction of Lcn2 synthesis over time was observed after treatment with cisplatin in the outer and inner medulla and an increase of its reabsorption by the proximal tubule cells (Figure 1A and supplementary Figure 1B). This effect was due to a transcriptional effect since Lcn2 mRNA expression increased progressively from D0 to D2 to reach a peak is maintained up to D4 (Figure 1B). This increasing expression was confirmed at the protein level by western-blot.

**Figure 1:**
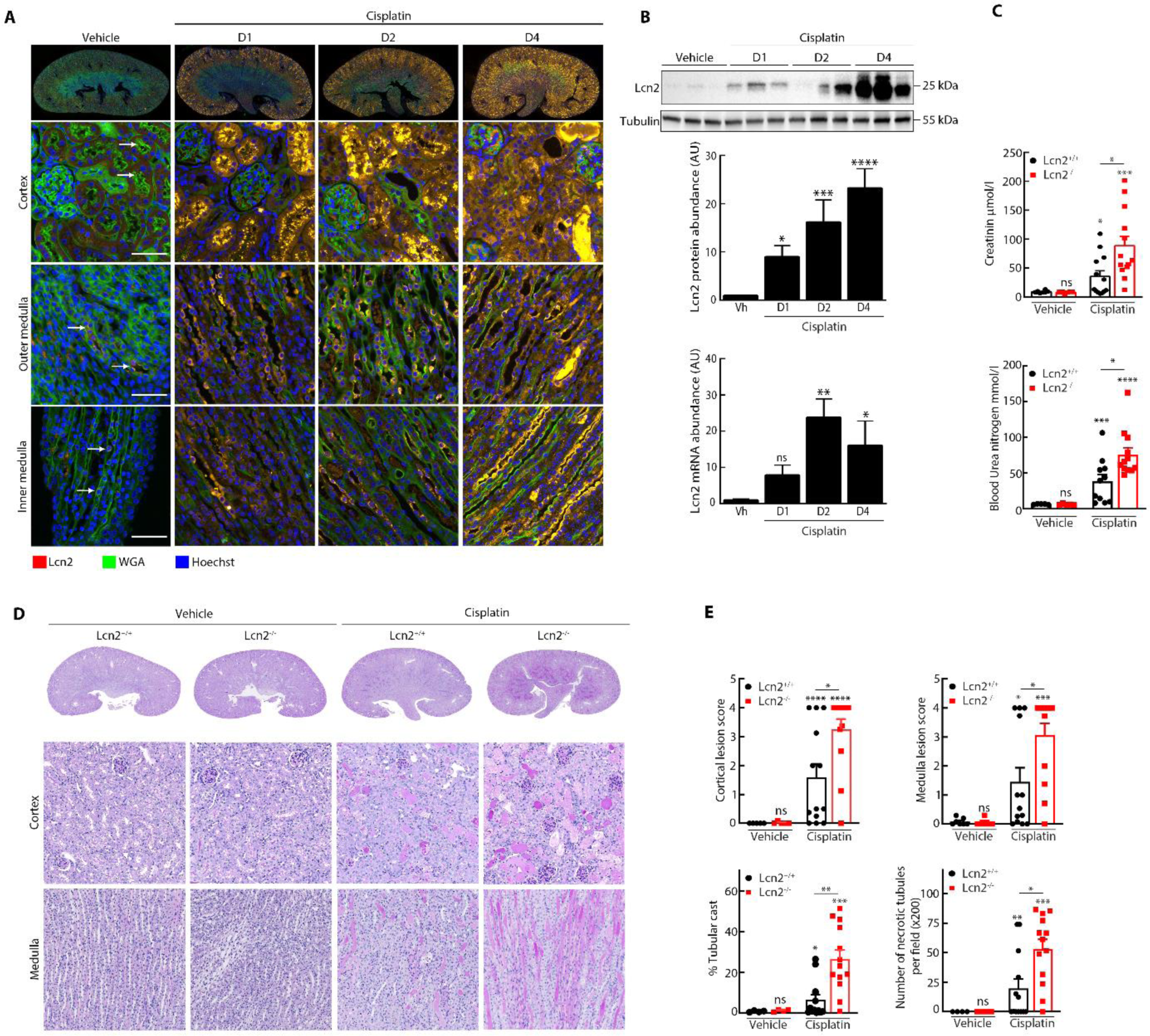
Invalidation of Lcn2 expression *in vivo* worsen cisplatin-induced kidney lesions. **(A)** Immunofluorescence on kidney sections showing Lcn2 (red), WGA (green) and nuclei counterstained with Hoechst in vehicle-treated *WT* (n=3) and cisplatin-treated *WT* (n=3) mice at day 1-2 or 4-. Original magnification x400. Scale Bar 50μM. **(B)** Representative Western-blot (top panel) and quantifications (bottom panel) of Lcn2 protein abundance and mRNA expression in *WT* mice 1 day, 2 days or 4 days after Cisplatin or Vehicle injection (n=12-8-11 mice/group respectively). **(C)** Evaluation of kidney function by measurement of plasma creatinine and blood urea nitrogen in vehicle-treated *Lcn2^+/+^* and *Lcn2^−/−^* (n=8) and cisplatin-treated *Lcn2^+/+^* and *Lcn2^−/−^* (n=13) mice. **(D)** Tubular morphology of renal cortex, (periodic acid–Schiff (PAS) staining) and renal medulla in vehicle-treated *Lcn2^+/+^* (n=4) and *Lcn2^−/−^* (n=2) and cisplatin-treated *Lcn2^+/+^*and *Lcn2^−/−^* (n=13) mice. Original magnification x200. **(E)** Quantification of histological damage by measuring the medulla and cortical lesions and the percentage of tubular cast and the number of necrotic tubules formation in vehicle-treated *Lcn2^+/+^* (n=4) and *Lcn2^−/−^*(n=2) and cisplatin-treated *Lcn2^+/+^* and *Lcn2*^−/−^ (n=13) mice. **(C-E)** Each dot represents one individual mouse. Data are means ± SEM. *p < 0.05; **p < 0.01; ***p < 0.001 versus Vehicle and *p < 0.05; **p < 0.01 *Lcn2^+/+^*; ***p < 0.001 versus *Lcn2^−/−^*, as determined by ANOVA followed by the Tukey– Kramer test.

To investigate the impact of Lcn2 expression on the outcome of CDDP accumulation in the kidney, *Lcn2^+/+^* and *Lcn2^−/−^*mice were subjected to a CDDP injection and were euthanized at different time points after injection. First, Serum creatinine and Blood urea nitrogen level in *Lcn2^−/−^* mice were significantly higher than in *Lcn2^+/+^* mice (Figure 1C and supplementary Figure 1D), indicating that Lcn2 tubular expression confers resistance to Cisplatin-induced AKI. Accordingly, PAS-stained histopathology revealed an increase of global lesions in medulla and cortical resulting in a significant increase of the percentage of tubular cast and the number of necrotic tubules per field in *Lcn2*^−/−^ kidneys compare to the *Lcn2^+/+^* (Figure 1D-E and supplementary figure 1E). The increased tubular injury observed in *Lcn2^−/−^* correlated with a stronger response to CDDP in term of DNA damage as shown by γH2AX immuno-staining in tubular cells (supplementary Figure 1F-G). Hence, CDDP induced expression of Lcn2 in the outer and inner medulla seems to participate to the protection of this compartment from the adverse effect of CDDP.

### Lcn2 protects renal tubular cell from apoptosis induced by cisplatin

Renal tissue damage, characterized by tubular cell death, is a common histopathological feature of cisplatin nephrotoxicity. Both necrosis and apoptosis have been described to participate to the tubular injury onset (Lieberthal *et al*, 1996).Hence, we examined if CDDP induced lesions in Lcn2 KO mice were a consequence of tubular cell death regardless of the pathway or if a specific cell death was favored by Lcn2 absence. Considering this, we first showed an overall increase of tubular cell death in Lcn2 KO mice compare to WT mice after CDDP injection using DNA fragmentation detection assay (TUNEL, TdT-mediated dUTP nick end labelling) (Figure 2A). To discriminate between apoptosis and necrosis, we then monitored respectively the cleavage of caspase 3 (CC3, apoptosis marker) and the induction of RIPK3 (necrosis marker) in our mice model. We observed (Figure 2B-C) that while CC3 cleavage was markedly increase in *Lcn2^−/−^* CDDP-treated mice, RIPK3 increase was comparable between WT and KO Lcn2 mice 4 days after injection. Furthermore, analysis of Bcl-2 (anti-apoptotic) and Bax (pro-apoptotic) ratio confirmed that the increase of apoptosis in *Lcn2^−/−^* mice might be responsible for the increase cell death in Lcn2 KO mice (Figure 2B).

**Figure 2:**
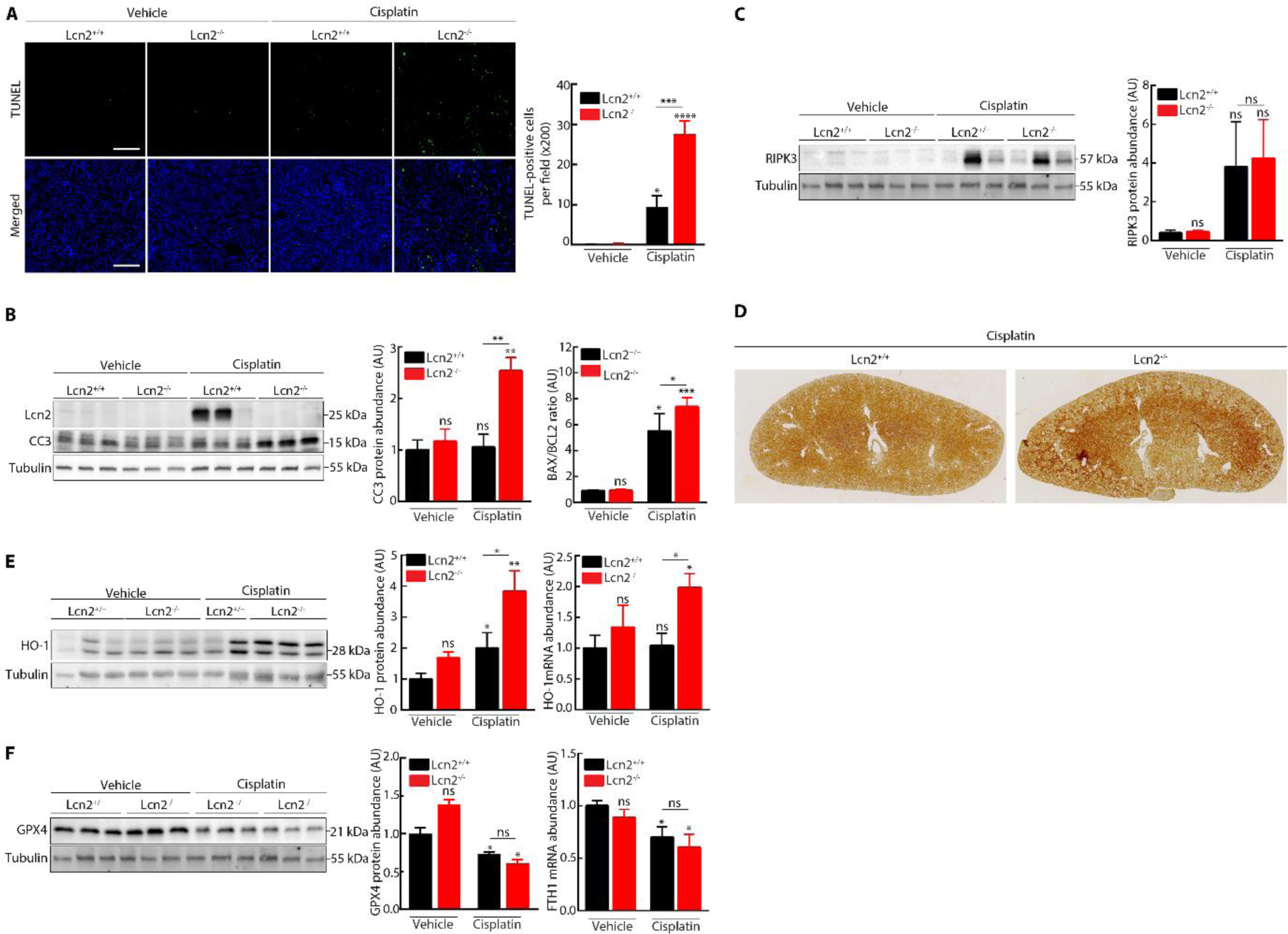
Lcn2 protects from tubular cell apoptosis induced by cisplatin. **(A)** Tubular cell apoptosis evaluated using TUNEL (left panels; scale bar 100 μm) and quantification (right panel) in vehicle-treated *Lcn2^+/+^*and *Lcn2^−/−^* (n=3) and cisplatin-treated *Lcn2^+/+^* and *Lcn2^−/−^* (n=3) mice. **(B)** Representative Western-blot (left panel) and quantifications (right panel) of CC3 protein abundance and mRNA expression of *BAX/BCL-2* ratio in *Lcn2^+/+^* or *Lcn2^−/−^* Vehicle and Cisplatin kidneys (n=10 mice/group). **(C)** Representative Western-blot (left panel) of necroptosis marker *RIPK3* and quantification (right panel) in *Lcn2^+/+^* or *Lcn2^−/−^* Vehicle and Cisplatin kidneys (n=6 mice/group). **(D)** Immunohistochemistry on kidney sections showing 4-HNE staining in cisplatin-treated *Lcn2^+/+^* and *Lcn2^−/−^*mice (D4). **(E)-(F)** Representative Western-blot and quantification and mRNA expression of oxidative stress marker *HO-1* (left panels) and ferroptosis markers *GPX4 and FTH1* (right panels) in *Lcn2^+/+^*or *Lcn2^−/−^* Vehicle and Cisplatin kidneys (n=6 mice/group). Data are means ± SEM. *p < 0.05; **p < 0.01; ***p < 0.001 versus Vehicle and *p < 0.05; **p < 0.01 *Lcn2^+/+^*versus *Lcn2^−/−^*, as determined by ANOVA followed by the Tukey–Kramer test.

It has been recently reported that ferroptosis, an iron-dependent oxidative cell death caused by ROS from the Fenton reaction and subsequent lipid peroxidation (Hu *et al*, 2019 ; Kuang *et al*, 2020) is involved in the pathogenicity of CDDP-induced AKI (Li *et al*, 2020). Considering the described effect of Lcn2 on iron homeostasis, we tested whether ferroptosis could participate to increased tubular lesion observed in Lcn2 KO mice. First, we determined the lipid peroxidation level in *Lcn2^+/+^* and *Lcn2^−/−^*mice treated with cisplatin. The staining of lipid peroxidation marker 4-hydroxynonenal (4-HNE) showed an increase in lipid peroxidation, mainly in the renal medulla, *Lcn2^−/−^* mice compare to *Lcn2^+/+^* (Figure 2D). Then, we monitored the expression of markers of oxidative stress and we showed a significantly stronger induction of HO-1 protein and mRNA abundance in *Lcn2^−/−^* CDDP-treated mice compare to *Lcn2^+/+^* (Figure 2E). However, we observed no difference for GPX4 and FTH1 mRNA, two markers of ferroptosis (Figure 2F). Hence, ferroptosis does not seem to be accountable for the effect of Lcn2 KO on CDDP-induced kidney injury. Together, these results indicate that Lcn2 ameliorated Cisplatin-induced acute kidney injury.

### Cisplatin regulates Lcn2 expression

To gain insight the mechanisms by which Lcn2 regulates the Cisplatin-induced apoptosis, we first studied *in vitro* the impact of CDDP on Lcn2 expression. To our knowledge, the direct effect of CDDP on Lcn2 mRNA and protein expression in a timely manner has never been described. Considering our *in vivo* results, we decided to use an immortalized inner medullary cell line (mIMCD-3) as a model. Constant exposure to CDDP in the culture media induced a dramatic decrease of Lcn2 mRNA and protein expression after 24h of stimulation (Figure 3A). Keeping in mind that *in vivo* exposure of kidney tubular cells to CDDP is described as been transitory (Chu *et al*, 2016), we tested the effect of a CDDP pulse (1h) chase (24h) exposure on Lcn2 expression. We observed that compared to continuous stimulation with cisplatin, transitory exposure to CDDP did not decrease Lcn2 expression but rather increased its mRNA and protein abundance between 24h and 72h of chase (Figure 3A-B). This direct effect of CDDP on Lcn2 expression in mIMCD-3 cells is strikingly comparable to the evolution of Lcn2 expression *in vivo* after CDDP injection.

**Figure 3:**
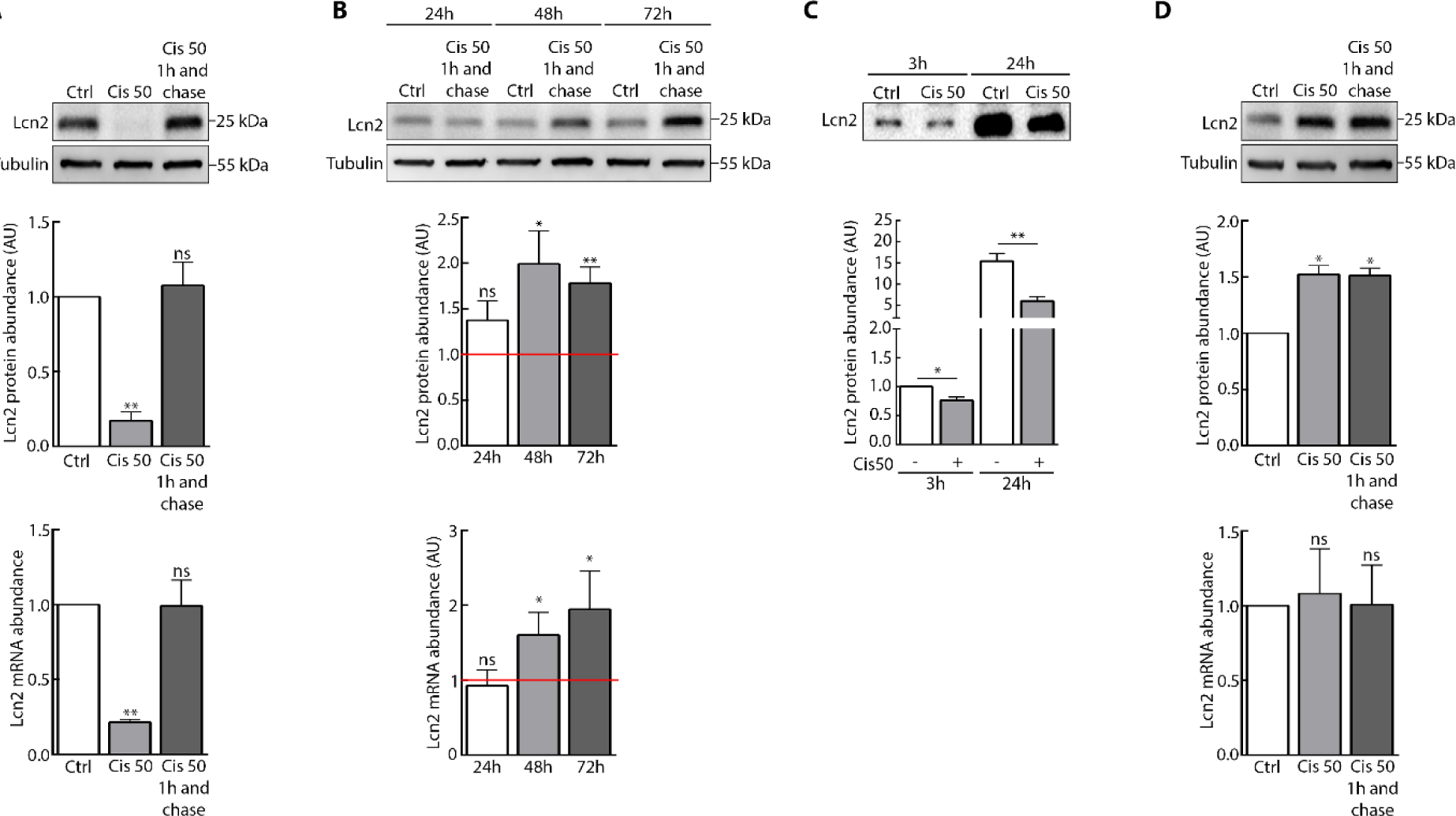
Cisplatin regulates Lcn2 expression. **(A)** WT mIMCD-3 cells were stimulated or not (Ctrl) with Cisplatin 50μM (Cis 50) for 24h or 1h (pulse) then chase for 23h. Representative Western-Blot (top panel) and quantification (bottom panels) of Lcn2 protein abundance and mRNA expression are shown (n=3). **(B)** WT mIMCD-3 cells were stimulated or not (Ctrl) with Cisplatin 50μM (Cis 50) for 1h (pulse) then chase for indicated time points. Representative Western-Blot (top panels) and quantification (bottom panels) of Lcn2 protein abundance and mRNA expression is shown (n=4). **(C)** WT mIMCD-3 cells were stimulated or not (Ctrl) with Cisplatin 50μM (Cis 50) for 3h or 24h. Representative Western-Blot (top panel) and quantification (bottom panel) of Lcn2 protein abundance in the supernatant are shown (n=3). **(D)** WT mIMCD-3 cells were stimulated or not (Ctrl) with Cisplatin 50μM (Cis 50) for 3h or 1h (pulse) then chase for 2h. Representative Western-Blot (top panel) and quantification (bottom panels) of Lcn2 protein abundance and mRNA expression are shown (n=4). Data are means ± SEM. *p < 0.05; **p < 0.01; ***p < 0.001; ****p<0.0001 versus controls, as determined by ANOVA followed by the Tukey– Kramer test.

In response to the formation of cisplatin-DNA crosslinks and adducts, a rapid DNA damage response which regulate multiple cellular responses is taking place. Considering that our inner medullary cells spontaneously expressed Lcn2, we wondered if CDDP could regulate Lcn2 independently of a transcriptional effect. Lcn2 being constitutively secreted while expressed, we monitored the presence of Lcn2 in the cell’s supernatant shortly after exposure du CDDP. Interestingly, CDDP decreased Lcn2 extracellular abundance compared to control condition as soon as 3h after treatment (Figure 3C). While the long-term exposure to CDDP decreased Lcn2 extracellular abundance most likely due to its effect on Lcn2 expression, our data showed that the decreased abundance of Lcn2 in the supernatant after 3h is most likely due to a blockage of its secretion since intracellular Lcn2 protein abundance increased at the same time (Figure 3D). The mechanism by which CDDP impaired Lcn2 secretion remain to be elucidate.

### The intracellular form of Lcn2 is responsible for the protective effect against Cisplatin

Our *in vivo* data demonstrated that CDDP induced more tubular apoptosis in Lcn2 KO mice than in WT. This observation led us to hypothesize that Lcn2 directly protects renal medulla tubular cells exposed to CDDP from apoptosis. To test this hypothesis, we first monitored the impact of Lcn2 KD *in vitro* on cell CDDP induced cell death. We observed that Lcn2 knockdown in mIMCD-3 cells expose to CDDP (constantly or pulse of 1h) induced more apoptosis than in control cells (Figure 4A, supplementary figure 2A). The effect of Lcn2 expression on CDDP-induced apoptosis was confirm in Lcn2-KO cells or Lcn2-overexpressing Lcn2 (supplementary figure 2B-C). Interestingly, we observed that LDH release was significantly increase in both scramble and sh-Lcn2 cells after cisplatin treatment (Figure 4B). Further experiment demonstrated that LDH release was mostly due to ferroptosis induction than necrosis since GPX4 decreased in both scramble and sh-Lcn2 cells after CDDP treatment while no effect was observed on RIPK3 abundance (Figure 4C). Accordingly, inhibitors of ferroptosis (Ferrostatin-1 or Deferoxamine) had no effect on the CDDP-induced cleavage of caspase-3 (supplementary Figure 2D-E).

**Figure 4:**
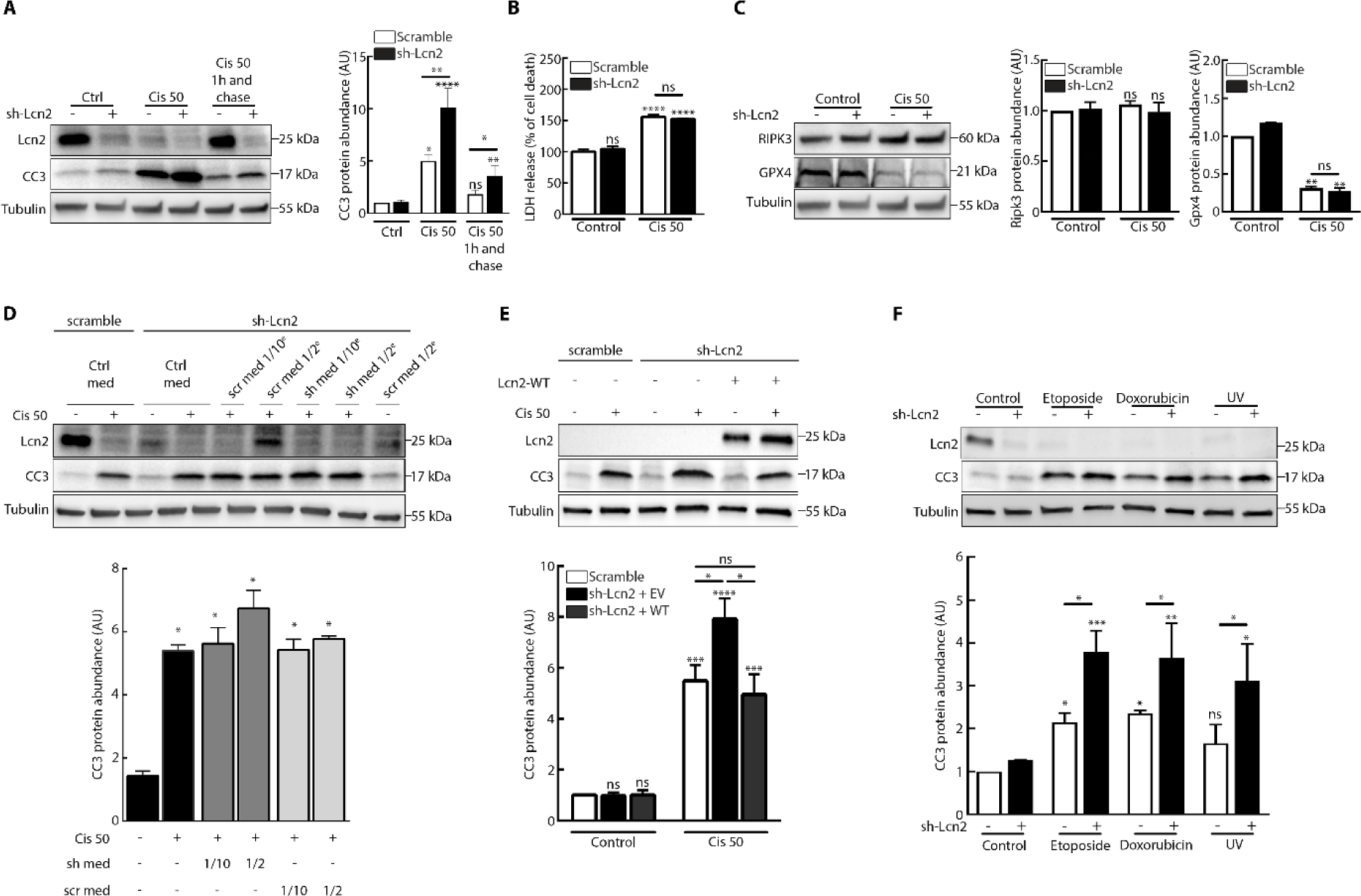
Lcn2 regulates cisplatin induced apoptosis. **(A)** Scramble and sh-Lcn2 mIMCD-3 cells were stimulated or not (Ctrl) with Cisplatin 50μM (Cis 50) for 24h or 1h (pulse) then chase for 23h. Representative Western-Blot (left panel) and quantification (right panel) of Cleaved-Caspase 3 (CC3) are shown (n=4). **(B)** Scramble and sh-Lcn2 mIMCD-3 cells were stimulated or not (Control) with Cisplatin 50μM (Cis 50) for 24h. Cell damage was measured by lactate dehydrogenase (LDH) release. **(C)** Scramble and sh-Lcn2 mIMCD-3 cells were stimulated or not (Control) with Cisplatin 50μM (Cis 50) for 24h. Representative Western-Blot of RIPK3 and GPX4 protein abundance (left panel) and RIPK3 mRNA abundance (right panel) are shown (n=2). **(D)** sh-Lcn2 mIMCD-3 cells were cultured with control media (Ctrl med) or with increased amount of mIMCD-3 Scramble (scr med) or sh-Lcn2 (sh med) cells conditioned media and stimulated (+) or not (−) with Cisplatin 50μM (Cis 50) for 24h. Western-Blot (top panel) show CC3 and Lcn2 protein abundance and quantification (bottom panel) show CC3 protein abundance (n=3). **(E)** Empty vector (−) or Lcn2-WT (+) transfected sh-Lcn2 mIMCD-3 cells were stimulated (+) or not (−) with Cisplatin 50μM (Cis 50) for 24 h. Representative Western Blot (top panel) of CC3 and Lcn2 protein and quantification (bottom panel) of CC3 abundance (n=4) are shown. **(F)** Scramble and sh-Lcn2 mIMCD-3 cells were stimulated or not (Control) with Etoposide 10μM, Doxorubicin 50μM or UVs 25mJ/cm^2^ for 24h. Representative Western-Blot of CC3 protein abundance are shown (n=3). Data are means ± SEM. *p < 0.05; **p < 0.01; ***p < 0.001; ****p<0.0001 versus control, *p < 0.05; **p < 0.01 scramble vs sh-Lcn2, as determined by ANOVA followed by the Tukey–Kramer test.

We next wondered whether the secreted form of Lcn2 was responsible for the effect on apoptosis after Cisplatin stimulation. To address this question, we treated sh-Lcn2 mIMCD-3 cells with conditioned medium obtained from cells secreting Lcn2 and stimulated these cells with Cisplatin for 24h. We demonstrated that addition of extracellular Lcn2 in the culture medium of Lcn2 failed to rescue cell death to the control level (Figure 4D). These results were confirmed by direct treatment of sh-Lcn2 mIMCD-3 cells with recombinant Lcn2 and stimulated them for 24h with Cisplatin (supplementary Figure 2F). By contrast, rescuing Lcn2 expression by transfection of sh-Lcn2 mIMCD-3 cells with Lcn2 overexpressing plasmid did decrease cell death to the control level (Figure 4E).

Finally, we wondered if Lcn2 protective effect CDDP-induced cell death was specific or if Lcn2 inhibited apoptosis in a genotoxic stress context regardless of the agent. We then exposed cells WT of Lcn2-KD to etoposide, doxorubicin or UV, all inducing DNA breaks and oxidative stress as CDDP. Hence, we demonstrated that Lcn2 expression protected cells from apoptosis induced by all these genoxotic agents (Figure 4F). Overall, these data strongly correlated with our *in vivo* observation, and demonstrated that Lcn2 expression protect IMCD cells from CDDP deleterious effect in a cell autonomous manner.

### The effect of Lcn2 on apoptosis is independent of the oxidative stress state

Cisplatin generates intracellular reactive oxygen species synthesis, which may be responsible for cell death and contributes to CDDP nephrotoxicity (Soni *et al*, 2018). We therefore wondered whether Lcn2 mediate CDDP resistance via the modulation the oxidative stress induced by it. First, we used MitoSOX probe to measure the generation of mitochondrial superoxide upon CDDP treatment. The same increase in the generation of superoxide after CDDP treatment was measured in both control and Lcn2-KD cells (Figure 5A). Furthermore, by measuring the abundance of the oxidative stress response protein HO-1, we observed no difference in the anti-oxidant response between control and Lcn2-KD cells (Figure 5B). Moreover, when adding N-Acetyl-Cysteine (NAC), ROS scavenger, to the medium, no effect was observed on the activation by CDDP of the apoptosis (Figure 5C-D).

**Figure 5:**
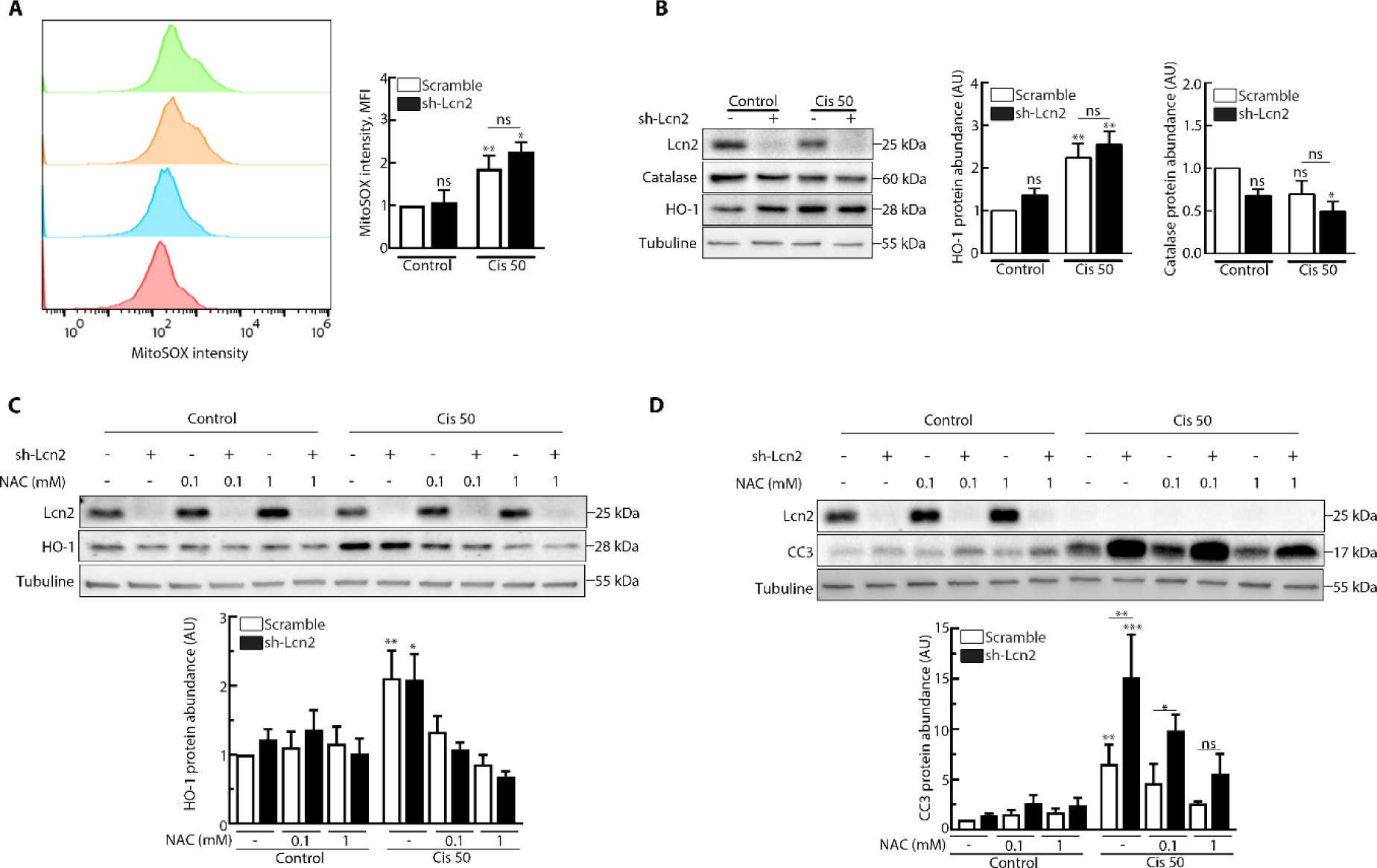
Impact of the blockage of oxidative stress on Lcn2 role in cisplatin-induced apoptosis. **(A)** Scramble and sh-Lcn2 mIMCD-3 cells were stimulated or not (Control) with Cisplatin 50μM (Cis 50) for 3h and treated with MitoSOX for 10 min. Flow cytometry was applied to analyze the level of ROS in cells and statistical analysis of ROS fluorescence is shown (n=3). **(B)** Scramble and sh-Lcn2 mIMCD-3 cells were stimulated or not (Control) with Cisplatin 50μM (Cis 50) for 3h. Representative western-Blot of HO-1 and Catalase, and quantification of HO-1 and Catalase (n=5) is shown. **(C)** Scramble and sh-Lcn2 mIMCD-3 cells were stimulated or not (Control) with Cisplatin 50μM (Cis 50) for 3h and treated with N-Acetylcysteine (NAC) 0,1mM. Representative western-Blot (left panel) of activation of antioxidant actor HO-1. Quantification of HO-1 (n=5) is shown. **(D)** Scramble and sh-Lcn2 mIMCD-3 cells were stimulated or not (Control) with Cisplatin 50μM (Cis 50) for 24h and treated with N-Acetylcysteine (NAC) 0,1mM. Representative western-Blot (left panel) of Cleaved-Caspase 3 (CC3), and quantification of CC3 (n=4) is shown. Data are means ± SEM. *p < 0.05; **p < 0.01; ***p < 0.001; ****p<0.0001 versus scramble control.

Thus, Lcn2 regulates the activation of DDR by genotoxic stresses independently of an effect on oxidative stress.

### Lcn2 regulates the DNA Damage Response pathway

To gain insight the mechanism by which Lcn2 regulates the cellular response to CDDP, we next monitored Lcn2 expression impact on the DNA Damage Response (DDR) pathway. Indeed, our data showing that Lcn2 expression can be regulated by CDDP shortly after exposure, we hypothesize that Lcn2 might regulate the DDR. Furthermore, has been reported that in renal cells, CDDP activates preferentially ATR, which in turn induces a p53 dependent apoptosis (Pabla *et al*, 2008). Hence, we monitored the DDR pathway in cells knockdown or not for Lcn2. We observed that in cells expressing Lcn2 ATM was activated by CDDP while in Lcn2 KD cells activated ATR (Figure 6A). Surprisingly, in both cell lines p53 was activated as well as H2AX. Nevertheless, Lcn2 KD cells had a greater induction of p21 expression. This difference in the DDR activation partners was not due to a difference in the damages triggered by CDDP since in the DNA breaks and the ROS generation observed shortly after CDDP exposure were comparable (Figure 6B and 5A). Considering the preferential activation of ATM in Lcn2 expressing cells, we next tested if ATM was required for the protective effect of Lcn2 on the CDDP-induced apoptosis. In the presence of ATM specific inhibitor KU-55933, CDDP triggered apoptosis in Lcn2 WT cells as well as in Lcn2-KD cells (Figure 6C). On the other hand, as previously described (Pabla *et al*, 2008), inhibition of p53 activity using pifithrin alpha (PFT) abrogated the difference of abundance of cleaved caspase 3 between Lcn2 WT and KD cells (Figure 6C).

**Figure 6:**
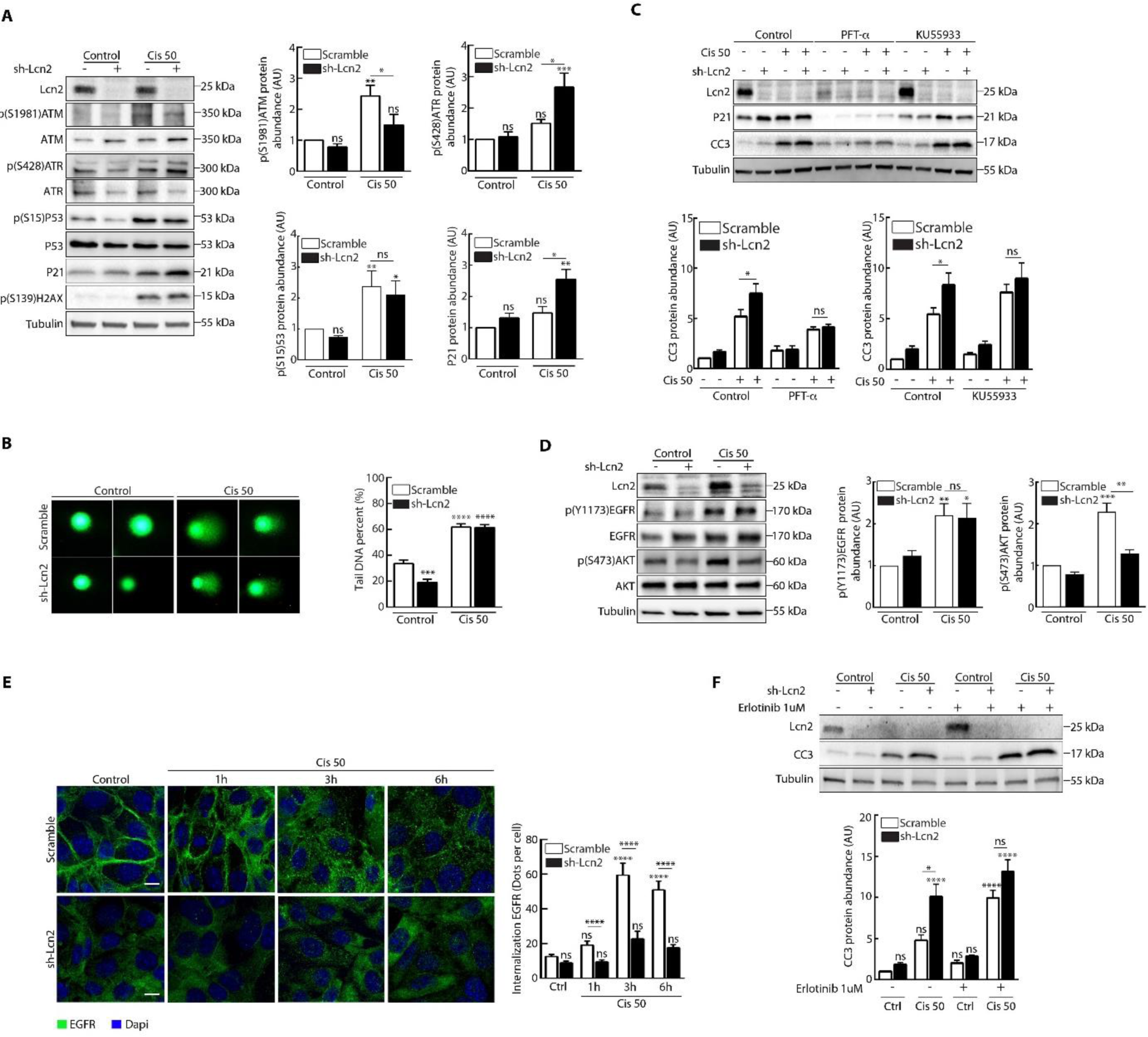
Lcn2 regulate cisplatin-induced apoptosis by activating EGFR and ATM pathway. **(A)** Scramble and sh-Lcn2 mIMCD-3 cells (Scramble) were stimulated or not (Control) with Cisplatin 50μM (Cis 50) for 3h. Representative western-Blot (left panel) of the DDR actors phosphorylation (p(S1981)ATM, p(S428)ATR, p(S15)P53, P21 and p(S139)H2AX respectively) and quantification of p(S1981)ATM, p(S428)ATR, p(S15)P53 and P21 (n = 6) is shown. **(B)** Scramble and sh-Lcn2 mIMCD-3 cells were stimulated or not (Control) with Cisplatin 50μM (Cis 50) for 3h. Comet Assay experiment shows DNA damage. The bright green dots represent the cellular nuclei area. Graph shows a quantification of the Tail DNA percent of comets (n=5). **(C)** Scramble and sh-Lcn2 mIMCD-3 cells were stimulated or not (Control) with Cisplatin 50μM (Cis 50) for 24h and treated with PFT 30μM or KU55933 10μM. Representative western-Blot (top panel) of Cleaved-Caspase 3 (CC3), and quantification of CC3 (n=4) is shown **(D)** Scramble and sh-Lcn2 mIMCD-3 cells were stimulated or not (Control) with Cisplatin 50μM (Cis 50) for 3h. Representative western-Blot (left panel) of EGFR and AKT phosphorylation (p(Y1173)EGFR and p(S473)AKT respectively) and quantification of p(Y1173)EGFR (n=6) and p(S473)AKT (n=3) is shown. **(E)** Scramble and sh-Lcn2 mIMCD-3 cells were stimulated with Cisplatin 50μM (Cis 50) for 1, 3 or 6h. Immunofluorescence shows EGFR (green) and nuclei counterstained with DAPI. Scale bar, 20μm. Graph shows a quantification of EGFR dots per cell (n=3). **(F)** Scramble and sh-Lcn2 mIMCD-3 cells were stimulated with Cisplatin 50μM (Cis 50) for 24h and treated with erlotinib 10μM. Representative western-Blot of Cleaved-Caspase 3 and quantification (n=3) is shown.

Finally, we decided to uncover the mechanism by which Lcn2 regulated the DDR in order to confer cellular resistance to CDDP-induced cell death. We have previously described that Lcn2 could regulate EGFR intracellular trafficking upon its activation and thus allowed a sustain activity of the EGFR pathway (Yammine *et al*, 2019). It has been reported that EGFR can be activated by CDDP and confer resistance to it (Tan *et al*, 2016). Furthermore, a study reported that EGFR can regulate ATM pathway activation (Lee *et al*, 2015a). We then hypothesis that Lcn2 regulates the DDR via the EGFR pathway. We first confirmed that EGFR was activated by CDDP (Figure 6D) but we did not see any difference in EGFR autophosphorylation between Lcn2 WT and KD cells. Nevertheless, we observed that Lcn2 WT cells had a robust activation of AKT compare to the Lcn2 KD cells. Intriguingly, it has been reported that CDDP can induce EGFR internalization and maintained perinuclear localization that allows the activation of AKT that leads to CDDP resistance (Winograd-Katz & Levitzki, 2006 ; Tomas *et al*, 2014). Having previously demonstrated than Lcn2 regulates EGFR trafficking, we followed EGFR intracellular journey upon CDDP exposure (Yammine *et al*, 2019). We observed that in cells expressing Lcn2 EGFR was internalized upon CDDP treatment and accumulated for up to 6h in vesicles (Figure 6E). This effect was not observed in Lcn2-KD cells. Then, to demonstrated that Lcn2 expression was blocking CDDP-induced cell apoptosis via its effect on EGFR activation, we tested the effect of EGFR inhibitor (erlotinib) in the presence of CDDP. Indeed, EGFR kinase activity inhibitor abrogated the effect of Lcn2 on apoptosis resistance observed after CDDP treatment (Figure 6F).

Together, the results indicated that Lcn2, via EGFR sustain activation, regulates the DDR pathway to stimulate pro-survival mechanisms in presence of cisplatin.

## DISCUSSION

Cisplatin induced nephrotoxicity represents the main limiting risk of cisplatin-based therapy (Miller *et al*, 2010). In acute kidney injury induced by cisplatin, or by a plethora of other causes, Lcn2 is strongly expressed shortly after the injury. Lcn2 presence in the blood and the urine is a biomarker of the renal parenchyma injury that anticipate the clinical diagnosis of the AKI (Mishra *et al*, 2003, 2004; Haase *et al*, 2011). Nevertheless, the role of Lcn2 induced expression in cisplatin-induced AKI remained elusive. In our study, we establish that Lcn2 renal tubular expression protects the parenchyma from the advert effect of cisplatin.

A recent study using Lcn2 reporter mice demonstrated that cisplatin triggers Lcn2 expression mostly in TAL and the CD in their medullary portion (Paragas *et al*, 2011) but not in the proximal tubule (PT). Furthermore, deep sequencing data obtained from microdissected rat kidney tubule segment (Lee *et al*, 2015) indicate that in steady state condition Lcn2 expression is detected in the TAL and the CD. In line with that, our data show that Lcn2 protein expression is detectable in the mTAL and the IMCD in control animals, and shortly after cisplatin injection, Lcn2 dramatically increases its expression in the same segments. Of note, Lcn2 protein was detected in control and cisplatin treated animals in the PT as a punctuate apical pattern most likely due to its reabsorption rather that its expression in the PT. Furthermore, we establish that Lcn2 expression in the kidney is protecting from tubular apoptosis rather than necrosis or ferroptosis. Paragas N et al, (Paragas *et al*, 2011) previously stated that Lcn2 expressing tubules were under cellular stress but without sign of apoptosis. Accordingly, our results point out Lcn2 as an induced stress protein that protects the tubular cells from cisplatin induced apoptosis. Using *in vitro* inner medulla collecting duct cells, we demonstrate that intracellular Lcn2 directly inhibits cisplatin-induced cells death, while the extracellular add no effect. We acknowledge the possibility that extracellular Lcn2 could regulate the cisplatin effect on other nephron segment, i.e. the proximal tubule. Overall, these results underline the fact that the renal medulla is an overlook compartment when it comes to the study of nephrotoxic effect of cisplatin. For instance, an elegant study by Miyaziaki et al (Miyazaki *et al*, 2019) recently reported that the collecting duct was a dominant source of ROS following AKI.

Previous studies established that Lcn2 expression protected cancer cells from cisplatin treatment (Miyamoto *et al*, 2016b; Rahimi *et al*, 2019; Huang *et al*, 2019). Nevertheless, the mechanisms driven by Lcn2 in order to promote cell survival after genotoxic exposure were unknown. Our study demonstrate that Lcn2 participates to the activation of EGFR pathway upon cisplatin exposure and EGFR pro-survival effect. These results confirm our previous report showing the role of Lcn2 on EGFR intracellular trafficking upon its activation by its ligand TGF-α (Yammine *et al*, 2019). It was already known that cisplatin can induce EGFR activation in a ligand-independent manner and its trafficking into the multivesicular bodies where it will be maintained and activates pro-survival pathways (Tan *et al*, 2016). Here we show that Lcn2 knockdown in IMCD cells abrogates EGFR intracellular trafficking and maintenance in the perinuclear region. This correspond to a loss of the EGFR dependent AKT activation observer in the WT cells. Further experiment demonstrate that EGFR is instrumental in the pro-survival effect of Lcn2 since blockage of EGFR tyrosine activity induces cell apoptosis. The mechanisms by which Lcn2 participates to the maintenance of EGFR in the multivesicular bodies are yet to be determine. Interestingly, interactome experiments have reported that Lcn2 can directly bind to protein such as Tumor susceptibility gene 101 protein (TGS101) and Sorting nexin-27 (SNX27) (Luck *et al*, 2020). This is of particular interest since we previously described that Lcn2 can localize in the cytosol and that TSG101 (and SNX27) is protein of the endosomal sorting complex required for transport (ESCRT) machinery showed to be required for the cisplatin induced intracellular retention of EGFR and its anti-apoptotic effect (Tomas *et al*, 2015).

We further demonstrate that Lcn2 expression favor ATM activation while Lcn2 knockdown induces an ATR activation upon cisplatin exposure. ATR pathway has been reported to be responsible for the p53 dependent cell apoptosis induction after cisplatin exposure in renal cells (Pabla *et al*, 2008). Interestingly, this study was performed on proximal tubular cells that to our knowledge do not express Lcn2. One could hypothesize that PT cells are more prone to ATR activation in absence of Lcn2 and that IMCD cells will activate ATM and its pro-survival effect. Indeed, our data demonstrate that ATM inhibition with a specific inhibitor induces apoptosis in WT cells while p53 inhibition decreases apoptosis in knockdown cells. The relationship between EGFR pathway activation remains to be clarified. Interestingly, it has been reported that EGFR can bind to ATM upon irradiation and phosphorylation ATM on its tyrosine 370 residue and hence regulates its activity (Lee *et al*, 2015a). Furthermore, Manohar et al. (Manohar *et al*, 2011) demonstrated that Chmp1A, a protein of the ESCRT system, can co-localize with ATM in MVBs where it activates ATM. It would be of great interest to characterize the putative role of Lcn2 in a MVB localized EGFR-ATM platform.

In summary, our study demonstrates for the first time that Lcn2 expression in medullary segment of the nephron is induced upon cisplatin exposure and protect the renal parenchyma from acute kidney injury. We establish that Lcn2 regulates an EGFR-ATM axis in IMCD cells that allows cell survival to genotoxic stress. In term of therapeutic, our study rise concerns about the use of Lcn2 as a therapeutic target to sensitize cancer cells to cisplatin since it may aggravate the nephrotoxicity of cisplatin and lead to chronic kidney disease.

## MATERIAL AND METHODS

### Animal procedures

All animal procedures performed in this study were approved by the Departmental Director of “Services Vétérinaires de la Préfecture de Police de Paris” and by the ethical committee of Université de Paris. Mice were housed in a specific pathogen-free facility, fed ad libitum and housed at constant ambient temperature in a 12-hour day/night cycle. Breeding and genotyping were done according to standard procedures.

A total of 15 eight weeks old male C57/BL6 strain mice invalidated (*Lcn2^−/−^*) or not (*Lcn2^+/+^*) for Lcn2 were injected with Nacl 0,9% only for vehicle mice. Mice in the experimental group were given a single intraperitoneal injection of cisplatin (Sigma-Aldrich, 479306), in the dose of 20 mg/kg. Mice were sacrificed 1, 2 or 4 days after Cisplatin injection. At sacrifice, kidneys were harvested for morphological, protein and mRNA studies.

### Cell culture and drug treatment

mIMCD-3 scramble and sh-Lcn2 cells were previously described (Yammine et al, 2019). Briefly viral transduction was performed with lentiviral vectors at a Multiplicity of Infection (MOI) of 5. Two days after transduction, sh-RNA Lcn2 transduced cells were selected in puromycin 2mg/ml for one week before efficacy was tested by qPCR and Western Blot. Cells were transfected with plasmids using Lipofectamine 2000 Reagent (Life Technologies) and Opti-MEM according to the manufacturer’s protocol. mIMCD-3 cells were cultured in 1:1 DMEM high-glucose pyruvate (41966052, GIBCO, Life Technologies): F-12 Nutrient Mixture (21765037, GIBCO, Life Technologies) containing 10% Fetal Bovine Serum (F7524, Sigma Aldrich) and 1% penicillin-streptomycin (15140122, Life technologies). All the cells were maintained at 37°C in a humidified atmosphere, containing 5% of CO2.

Cisplatin (5μM; Sigma) was dissolved in 0.9% NaCl. Erlotinib (10μM; Sigma) was dissolved in dimethyl sulfoxide (DMSO; Sigma). N-Acetylcysteine (NAC 0,1mM; Sigma) was dissolved in deionized water. Deforoxamine (DFO 50μm; Sigma) was dissolved in DMSO. Pifithrin-α was dissolved in DMSO (PFT 30μm; Santa cruz Biotechnology). KU55933 was dissolved in DMSO (10μm; Sigma). Ferrostatin-1 (Fer-1 1μm; Sigma) was dissolved in DMSO. For NAC, PFT, KU55933 and Erlotinib the treatment was added at confluency 30 min before Cisplatin stimulation.

### Plasmid constructs

pcDNA3.1-c-DYK and pcDNA3.1-mouse-Lcn2-c-DYK were purchased from Genscript.

### Transient transfections

Cells were transfected with plasmids using Lipofectamine 2000 Reagent (Life Technologies) and Opti-MEM according to the manufacturer’s protocol. The transfection mixture was removed 24 h later and was replaced with a fresh medium. Subsequent experiments on cells started 24 h later.

### Western Blot Analysis

Cells were lysed in a buffer composed of 150 mM NaCl, 0.5% Sodium Deoxycholate, 0.1% SDS, 50 mM Tris-HCl pH 8.0 with protease and protein phosphatase inhibitors tablets (Thermo Fisher Scientific, A32959). After centrifugation for 15 min, the total protein concentration was measured in cell extracts according to the Bradford method using the BCA Protein Assay kit (Pierce), then denatured and reduced in a Laemmli-b-mercaptoethanol mixture at 95°C for 5 min. Then an equal amount of proteins were separated on a polyacrylamide gel with pre-cast gradient (Mini-PROTEAN® Precast Gels, acrylamide 4% −20%, Bio-Rad) and transferred to nitrocellulose membrane (Trans-Blot® Turbo ™ Midi Nitrocellulose Transfer Packs, Bio-Rad) using the Trans-Blot® Turbo ™ (Bio-Rad). Non-specific binding sites were blocked by incubation with a blocking solution (3% BSA in TBS-Tween 0.1%) for 30 min. Subsequently, membranes were incubated with the primary antibodies (Listed in Table 1) overnight at 4°C. After washing, membranes were incubated with Alexa Fluor 647, Alexa Fluor 488 donkey anti-specified specie (Molecular Probes, Life Technologies) or with HRP conjugated secondary antibodies for 30 min. The immunoreactive proteins were detected by chemiluminescence (ECL kit; Amersham). The membranes were revealed using the ChemiDoc Imaging System Bio-Rad and the signals quantified with the Image Lab software (Bio-Rad).

**TABLE 1:**
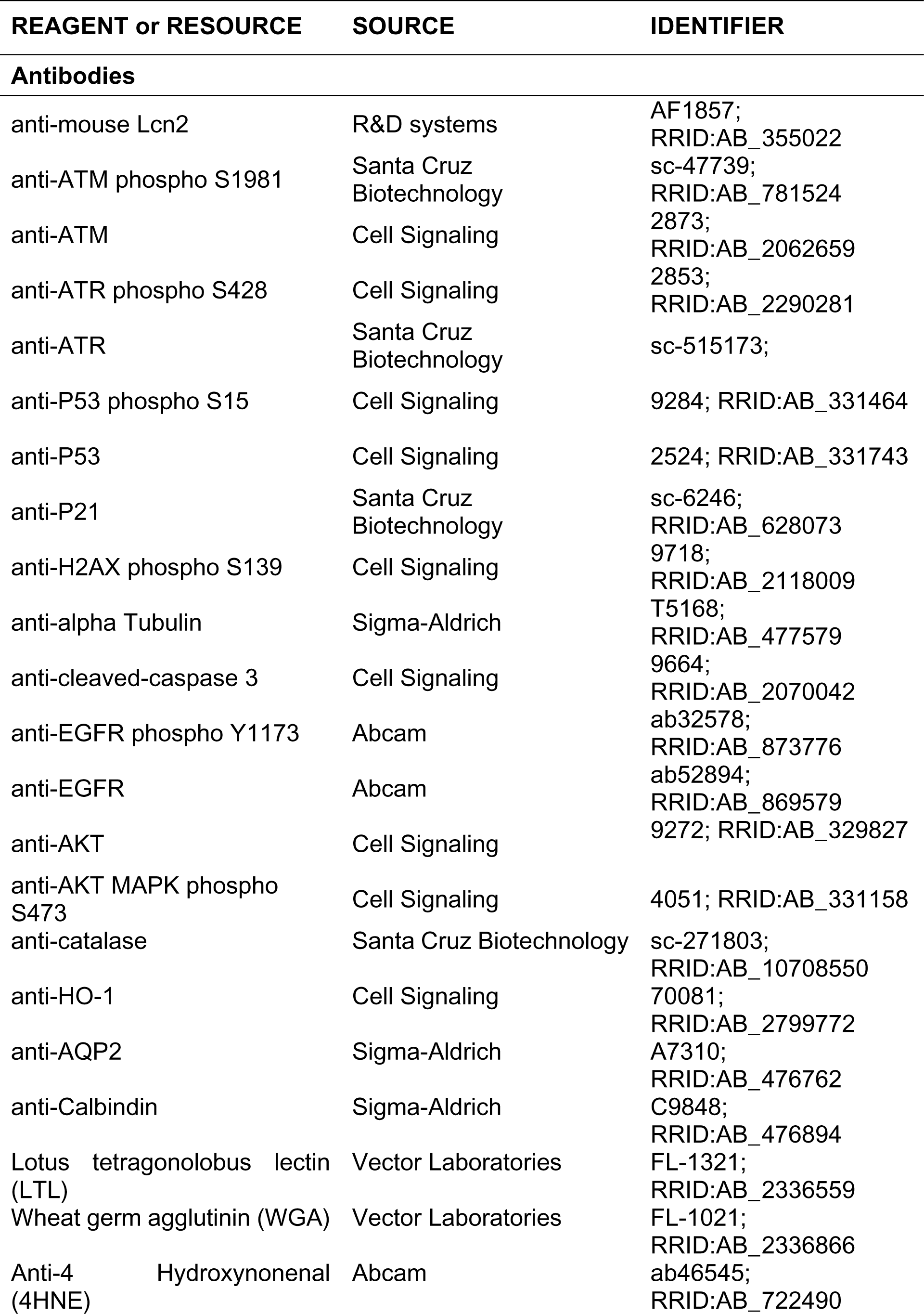
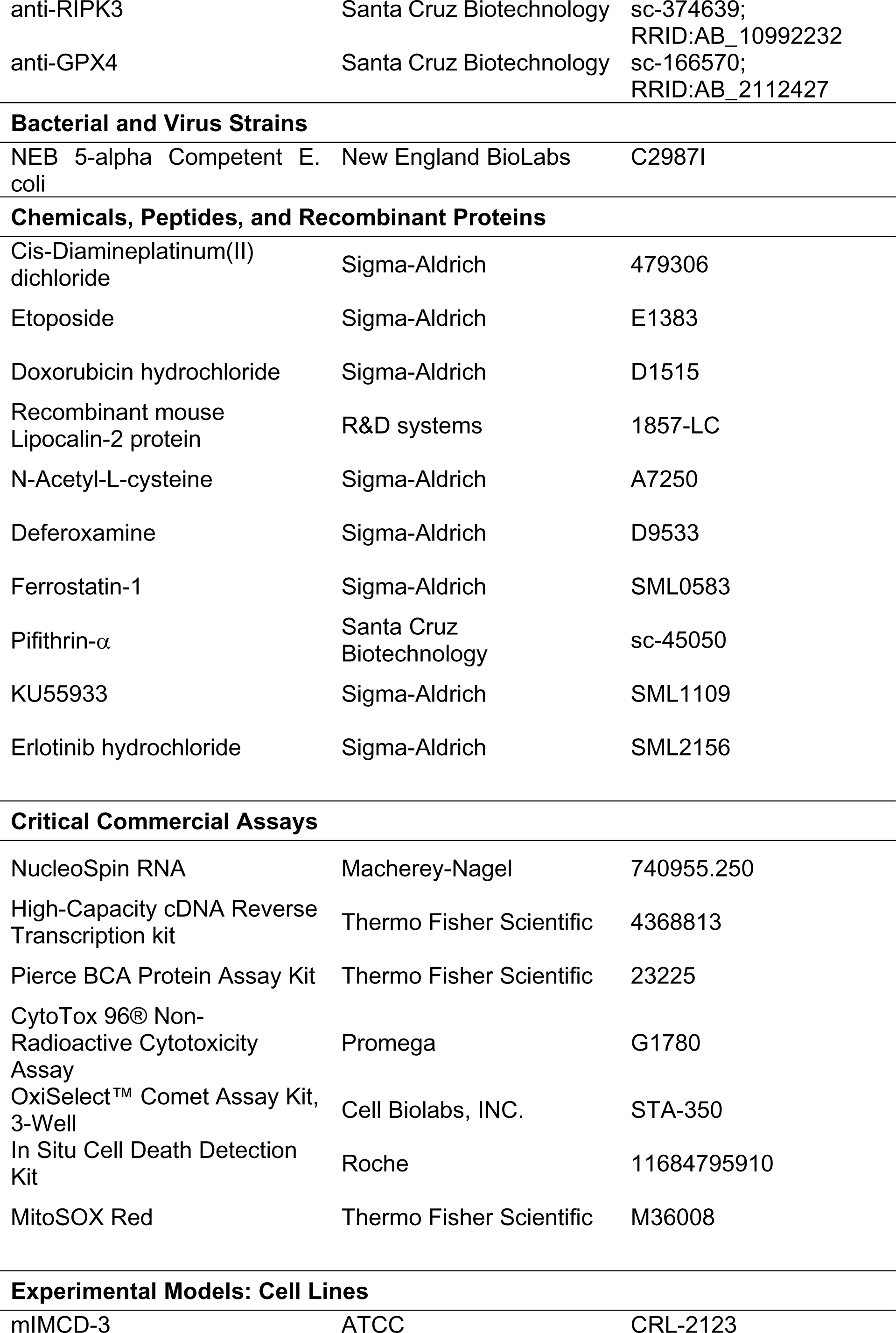
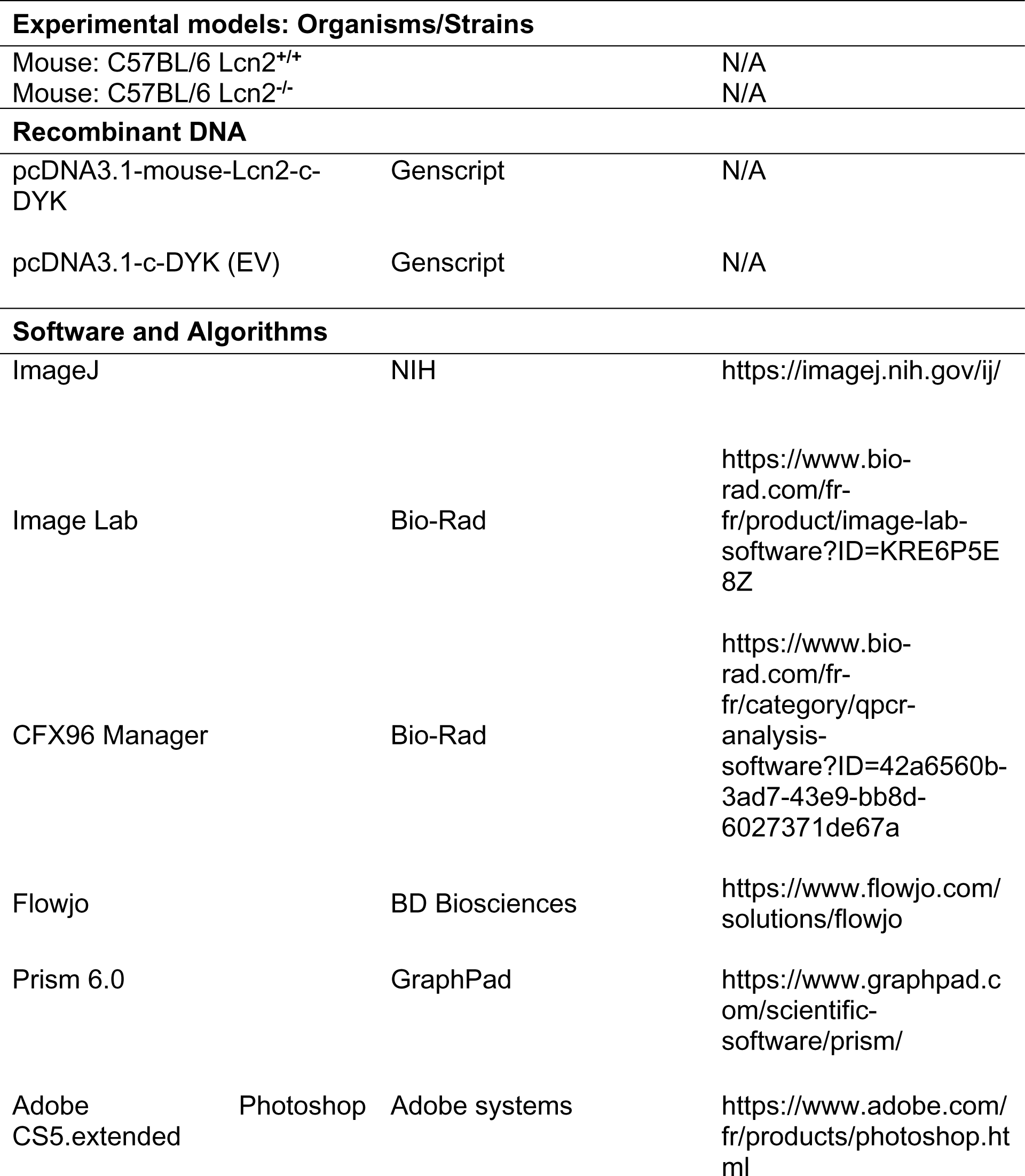
Key resources.

### Flow cytometry detection of reactive oxygen species (ROS)

The intracellular ROS levels in mIMCD-3 scramble and sh-Lcn2 cells were monitored using MitoSOX (ThermoFischer Scientific M36008) and examined by flow cytometry. Briefly, mIMCD-3 scramble and sh-Lcn2 cells at a seeding density of 2 × 10^5^ cells/well, were seeded in a six well plate. At confluency, cells were trypsinized and washed twice with PBS, and then treated with MitoSOX 5μM. After being incubated at 37°C for 10 min, the levels of ROS were analyzed by FACS using a SP6800 flow cytometer (Sony) at an excitation 510 nm and emission 580 nm. The data were processed and analyzed using FlowJo software.

### Comet assay (single-cell gel electrophoresis assay)

Comet assay were performed according to the manufacturer’s recommendations (OxiSelect Comet Assay Kit, Cell Biolabs, Inc). Briefly, following treatment with the Cisplatin for 3h, the cells were harvested, washed and re-suspended in PBS. 3×10^−5^ of cell suspensions were mixed with 90µl 0.5% agarose at 37°C. 75μl of the cell-agarose mix were added to Comet slides. The slides were transferred at 4°C in the dark for 15 min to accelerate gelation. When the agar was solidified, embedded cells were submerged in fresh pre-chilled lysis buffer at 4°C during 45min. Lysis Buffer was replaced with cold alkaline Solution and placed at 4°C for 30min in the dark. Electrophoresis experiment (21V for 23 min at 4°C) was performed in a horizontal electrophoresis chamber filled with cold Alkaline Electrophoresis Solution. The slides were rinsed 3 times in cold distilled-water for 2 min and then placed in 70% ethanol for 5 min and allowed to dry. Once the slides are completely dry, 100 μL/well of diluted Vista Green DNA Dye were added for 15min at room temperature. Comets were visualized with a fluorescence microscope (Nikon Eclipse E800) at ×100 magnification using a ×10 objective. Images were analyzed with the OpenComet software. A total of 50 cells were analyzed per sample. Tail Moment, which indicates damaged DNA, was calculated for each cell.

### mRNA analysis

mRNAs were quantified in mouse kidneys and cultured cells by quantitative RT-PCR RNA extraction performed as described in the manufacturer’s protocol (NucleoSpin RNA, Macherey-Nagel). cDNA was synthesized with the High-Capacity cDNA Reverse Transcription kit (ThermoFisher Scientific), Quantitative PCR was performed with iTaqUniversal SYBR Green Supermix (Biorad) using primers listed Table 2. RPL13 and HPRT were used as the normalization controls.

**TABLE 2:**
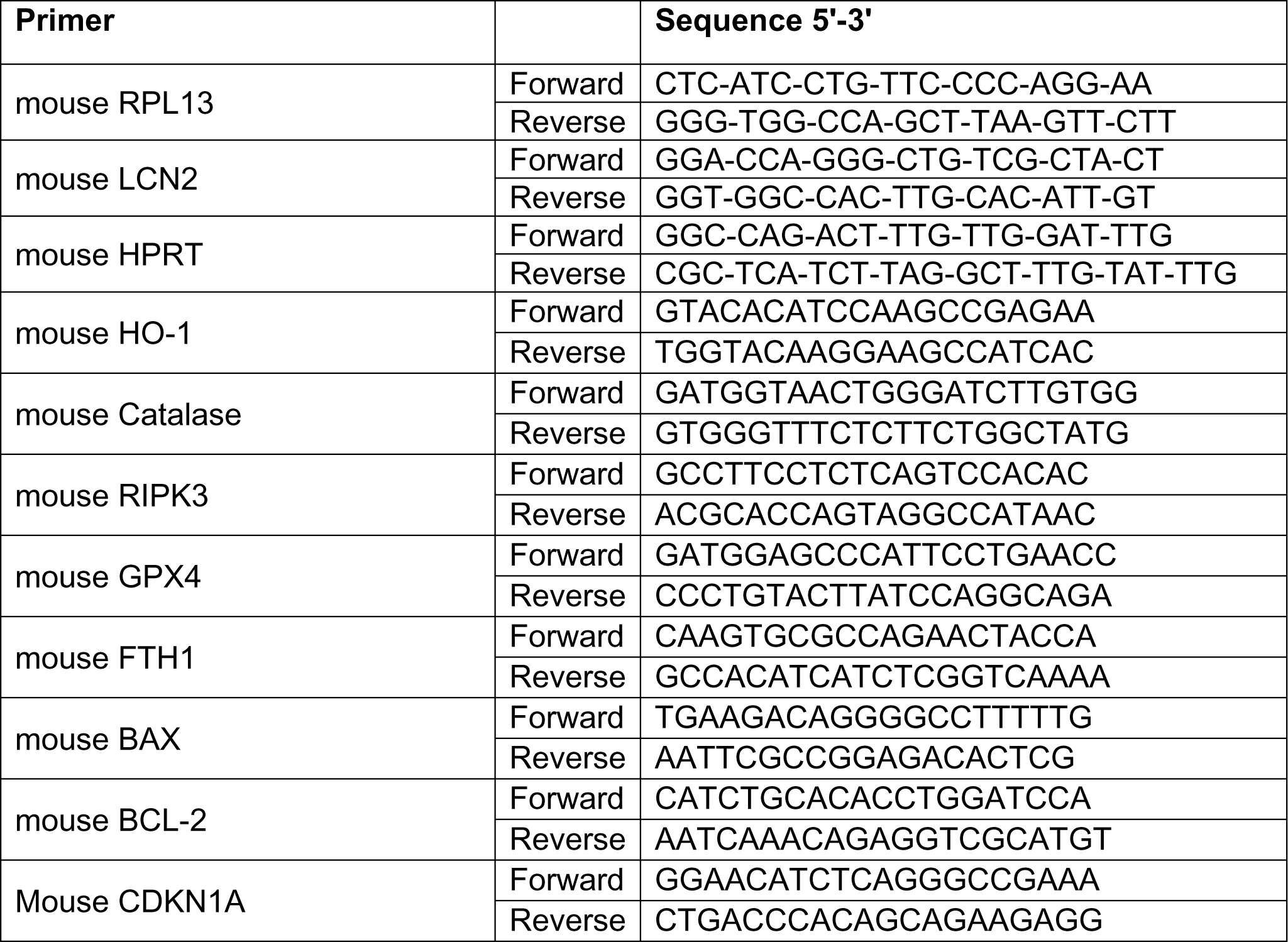
List of primers used in this study.

### Immunofluorescence

#### In vitro

The cells were plated on coverslip and then stimulated at confluence at the times indicated with 50 μM Cisplatin. The cells were washed with PBS, fixed in 4% paraformaldehyde for 15 min. Then cells were permeabilized in 0.1% Triton in PBS for 5 minutes and blocked in 3% BSA - 0,1% Triton in PBS for 30 min at room temperature. They were then incubated overnight at 4°C in blocking buffer with the following primary antibodies. This was followed by incubation with Alexa 488 or 555-conjugated secondary antibodies for 1 h at room temperature. Nuclei were counterstained with DAPI for 5 min. The slides were mounted using a Fluoromount™ aqueous mounting support (Sigma-Aldrich) and examined by fluorescence microscopy (Nikon Eclipse E800). The number of internalized dots per cell reflecting EGFR internalization have been quantified with ImageJ.

#### In vivo

The 4 μm sections of kidneys, included in paraffin, were subjected to an antigen retrieval in a citrate buffer pH 6.0 (ZUCD28; Zytomed) and heating in high pressure cooker for 15 minutes. Nonspecific binding was blocked with a buffer containing 3% BSA, 10% FBS in PBS for 30 minutes. Sections were incubated with primary antibodies, in blocking buffer overnight at 4°C. After several washes, the sections were incubated with Alexa-555, 488 or 647-conjugated secondary antibodies for 1h at room temperature and the nuclei were counterstained with Hoechst. The slides were mounted using a Fluoromount™ aqueous mounting support (Sigma-Aldrich) and examined by confocal microscopy (Zeiss LSM700), except for figure 1A and supplementary figure 1A where images were acquired using NanoZoomer 2.0 HT (Hamamatsu).

### Morphological Analysis

Mouse kidneys were fixed in 4% paraformaldehyde, paraffin embedded and 4 μm sections were stained with periodic acid–Schiff (PAS). Stained sections were imaged using an NanoZoomer 2.0 HT (Hamamatsu). The degree of tubular lesions was evaluated using semiquantitative score methodology as previously described, with minor modifications in the evaluation of kidney lesions (0 = no lesions, 1 = involvement of < 25% of the cortex or medulla, 2 = involvement of 25-50% of the cortex or medulla, 3 = involvement of 50-75% of the cortex or medulla, 4 = involvement of 75-100% of the cortex or medulla. The number of necrotic tubules and tubular cast per field was measured with ImageJ software from PAS stained x20 kidney images.

### Kidney function

For mouse samples, plasma urea nitrogen and creatinine levels were determined using a Konelab 20i analyser (Thermo Fisher Scientific; Instrumentation Laboratory).

### Immunohistochemistry

4µm sections of paraffin-embedded kidneys were submitted to avidin/biotin blocking (Vector, SP-2001). Sections were incubated with primary antibody followed by biotinylated antibody. Biotinylated antibodies were detected using HRP-labelled streptavidin (Southern Biotech, 7100-05) at 1:500 and 3-3’-diamino-benzidine-tetrahydrochloride (Dako, K3468). Stained sections were imaged using an NanoZoomer 2.0HT (Hamamatsu).

### TUNEL assay

Apoptosis was measured by Terminal Deoxynucleotidyl Transferase-Mediated dUTP-Biotin Nick End Labeling (TUNEL) assay using In Situ Cell Death Detection kit (Roche) according to the manufacturer’s protocol. Detection of the apoptotic cells showing green fluorescence was performed by fluorescence microscopy. Nuclei were counterstained with DAPI. Ten random images of both the cortex and medulla region of the kidney were taken under 200x magnification using and quantified by ImageJ software (NIH). The number of TUNEL-positive cells was divided by the total number of cells in each image to obtain the percentage of TUNEL-positive cells.

### Quantification and statistical analysis

All experiments were repeated at least three times. The exact number of replications for each experiment is detailed in text and figures. Data are expressed as means ± SEM. Quantitative data were analyzed using Prism 6.0 software (GraphPad). Groups were compared with an analysis of variance (ANOVA) or a Student’s t test, P value < 0.05 was considered as significant.

## Acknowledgements

We are grateful to Sylvie Fabrega and to all the persons working in the viral vectors and Gene transfer facility. We are grateful to Meriem Garfa-Traore and to all the persons working in the Cellular Imaging facility. We are grateful to Sophie Berissi and to all the persons working in the Morphology and LEAT facilities for technical assistance. This work was supported by INSERM, Université Paris Descartes and Agence Nationale Recherche.

## Footnotes

### Author contributions

**A.Z.** designed and performed the experiments, analysed the data and wrote the paper. **E.M, L.Y**. performed *in vitro* studies. **C.N** performed *in vivo studies* **F.T.** provided the conceptual framework, designed the study and revised the manuscript. **M.G.** provided the conceptual framework and designed the study, supervised the project and wrote the paper.

---

**Supplementary figure 1:**
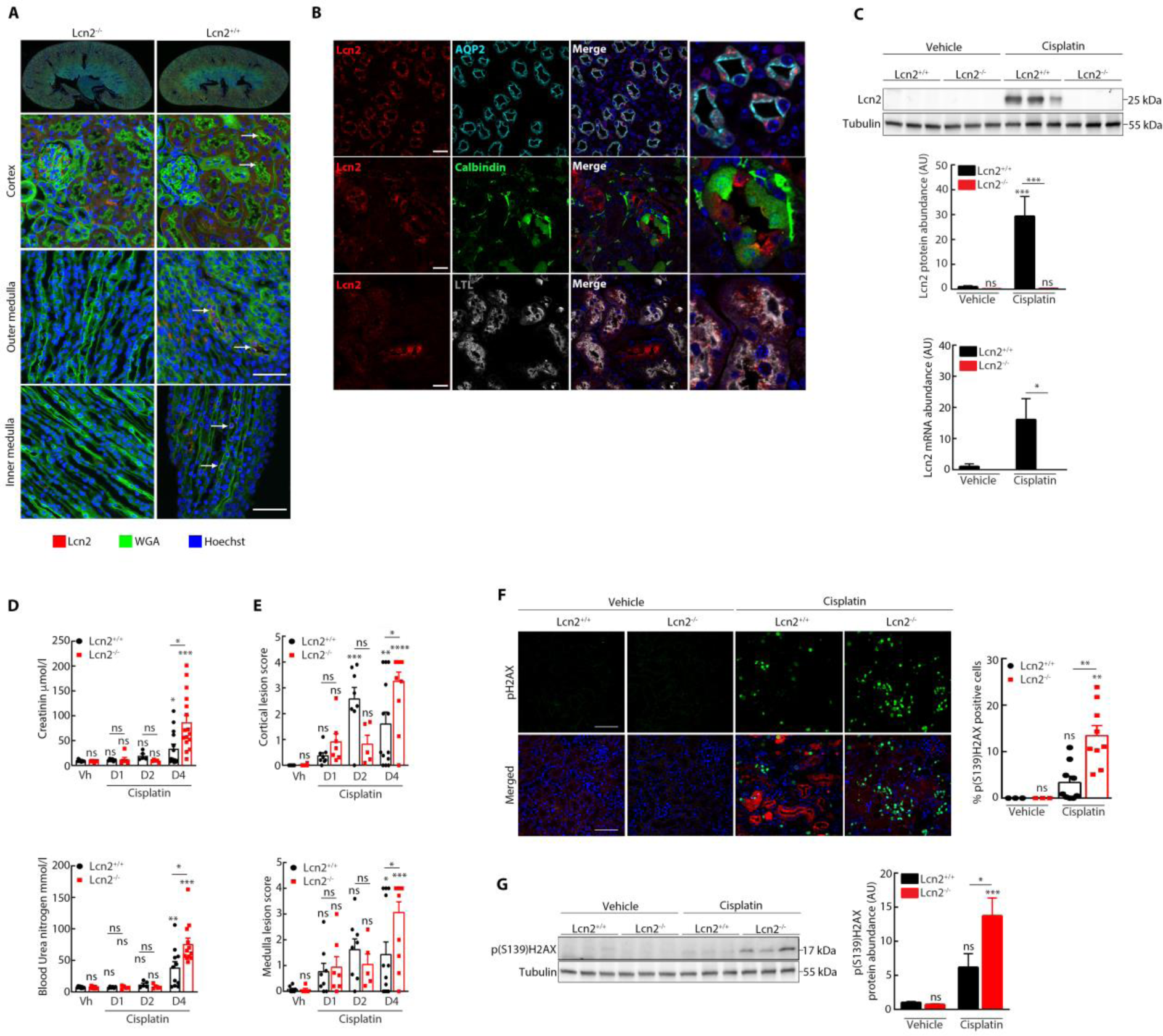
**(A)** Immunofluorescence on kidney sections showing Lcn2 (red), WGA (green) and nuclei counterstained with Hoechst in vehicle-treated *Lcn2^+/+^* and *Lcn2 ^−/−^* (n=3) mice. Original magnification x400. Scale Bar 50μM. **(B)** Immunofluorescence on kidney sections showing Lcn2 (red), AQP2 (cyan), Calbindin (green), LTL (white) and nuclei counterstained with Hoechst in cisplatin-treated *Lcn2^+/+^* and *Lcn2 ^−/−^* (n=3) mice at day 2. Original magnification x400. Scale Bar 20μM. **(C)** Representative Western-blot (top panel) and quantification (bottom panel) of Lcn2 protein and mRNA abundance at day 4 in *Lcn2^+/+^*or *Lcn2^−/−^* Vehicle and Cisplatin kidneys (n=6 mice/group). **(D)** Evaluation of the evolution of kidney function by measurement of plasma creatinine and blood urea nitrogen in *Lcn2^+/+^*and *Lcn2^−/−^* 1 day, 2 days or 4 days after Cisplatin or Vehicle injection. **(E)** Quantification of histological damage by measuring the medulla and cortical lesions and the percentage of tubular cast and the number of necrotic tubules formation in *Lcn2^+/+^* and *Lcn2^−/−^* 1 day, 2 days or 4 days after Cisplatin or Vehicle injection. **(F)** Immunofluorescence on kidney sections showing pH2AX (green), Lcn2 (red) and nuclei counterstained with Hoechst (left panels; scale bar 100 μm) and quantification (right panel) in vehicle-treated *Lcn2^+/+^* and *Lcn2^−/−^* (n=3) and cisplatin-treated *Lcn2^+/+^* and *Lcn2^−/−^* (n=9) mice. **(G)** Representative Western-blot (right panel) and quantification (left panel) of pH2AX protein abundance in *Lcn2^+/+^* or *Lcn2^−/−^* Vehicle and Cisplatin kidneys (n=13 mice/group). Data are means ± SEM. *p < 0.05; **p < 0.01; ***p < 0.001; ****p<0.0001 versus scramble control.

**Supplementary figure 2:**
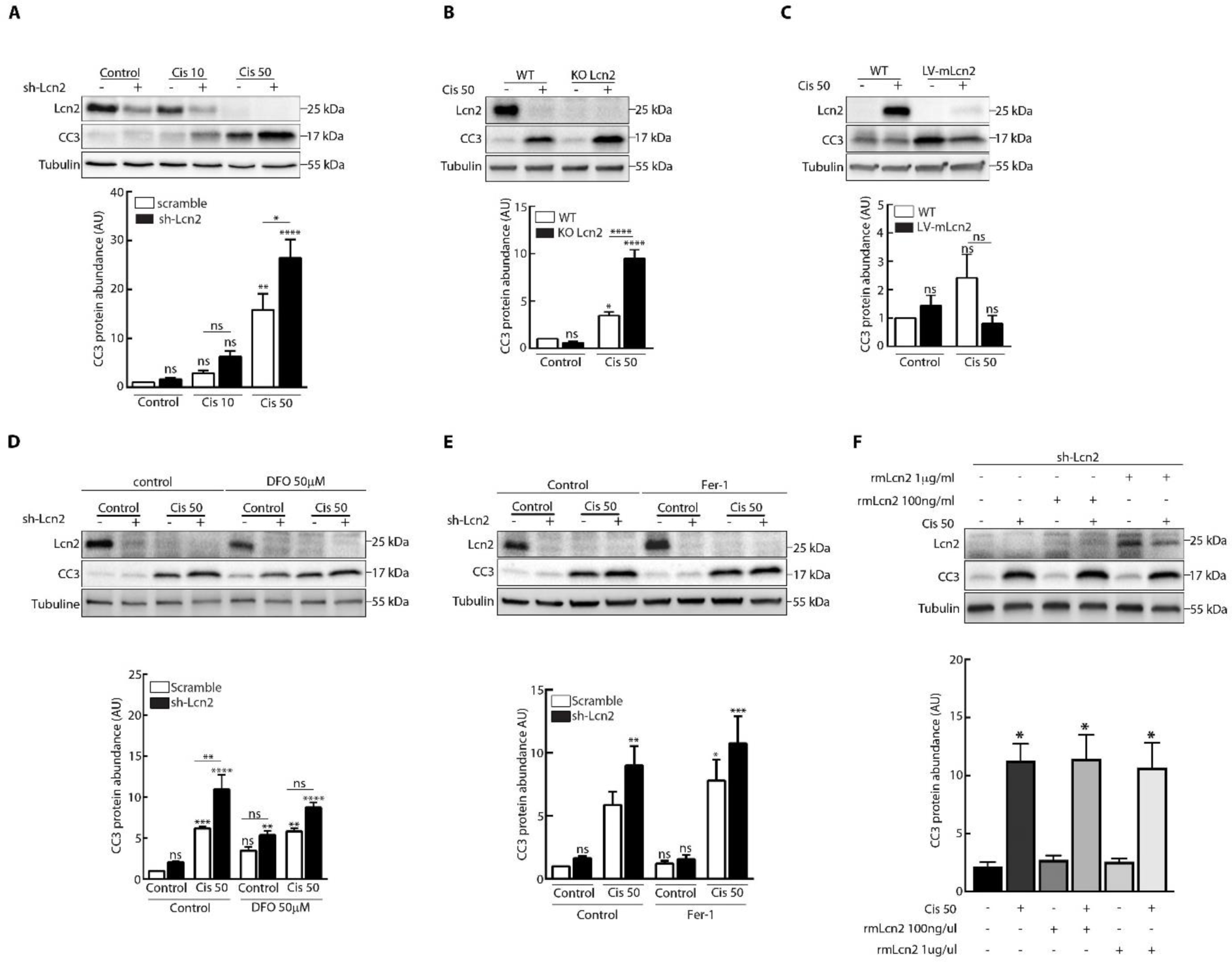
**(A)** Scramble and sh-Lcn2 mIMCD-3 cells cultivated on filter were stimulated or not (Control) with Cisplatin 10μM or 50μM (Cis 50) for 24h. Representative Western-Blot (top panel) and quantification (bottom panel) of Cleaved-Caspase 3 (CC3) are shown (n=4). **(B)** Scramble and KO-Lcn2 mIMCD-3 cells were stimulated or not (Control) with Cisplatin 50μM (Cis 50) for 24h. Representative Western-Blot (top panel) and quantification (bottom panel) of Cleaved-Caspase 3 (CC3) are shown (n=4). **(C)** WT and overexpressed (Lv-mLcn2) mIMCD-3 cells were stimulated or not (Control) with Cisplatin 50μM (Cis 50) for 24h. Representative Western-Blot (top panel) and quantification (bottom panel) of Cleaved-Caspase 3 (CC3) are shown (n=4). **(D)** Scramble and sh-Lcn2 mIMCD-3 cells were stimulated or not (Control) with Cisplatin 50μM (Cis 50) for 24h and treated with deferoxamine (DFO) 50μM. Representative western-Blot (top panel) of Cleaved-Caspase 3 (CC3), and quantification of CC3 (n = 4) is shown. **(E)** Scramble and sh-Lcn2 mIMCD-3 cells were stimulated or not (Control) with Cisplatin 50μM (Cis 50) for 24h and treated with ferrostatin-1 (Fer-1) 1μM. Representative western-Blot (top panel) of Cleaved-Caspase 3 (CC3), and quantification of CC3 (n=4) is shown. **(F)** sh-Lcn2 mIMCD-3 cells were stimulated or not with Cisplatin 50μM (Cis 50) for 24h with (+) or without (−) mouse recombinant Lcn2 (rmLcn2) at 100ng/ml or 1μg/ml. Representative Western-Blot (top panel) and quantification (bottom panel) of CC3 protein abundance is shown (n=3).

